# Targeting *Runx*1 protects against heart failure with preserved ejection fraction

**DOI:** 10.1101/2025.01.24.634831

**Authors:** Ali Ali Mohamed Elbassioni, Anmar A Raheem, Jian Song, Alexander S Johnston, Cara Trivett, Hong Lin, Haobo Zhang, Ashley Bradley, Erin Higgins, Leanne Mooney, Yen Chin Koay, Dylan O’Toole, Pawel Herzyk, Colin Nixon, Karen Blyth, John F O’Sullivan, Ninian N Lang, Colin Berry, Thomas Braun, Gabriele G Schiattarella, Mauro Giacca, Martin W McBride, Stuart A Nicklin, Ewan R Cameron, Christopher M Loughrey, Eilidh A MacDonald

**Author notes:** AAME and AAR, as well as. EAM and CML contributed equally to this work and as joint first/last authors can change the order of authorship for the purposes of curriculum vitae.

## Abstract

Heart failure with preserved ejection fraction (HFpEF) is a public health problem and an elusive illness for which there are few treatment options. HFpEF is a systemic condition with a broad phenotype including diastolic dysfunction, pulmonary oedema, exercise intolerance, and left ventricular (LV) hypertrophy, collectively resulting in enhanced morbidity and mortality. Master-regulator transcription factor RUNX1 has recently been identified as a mediator of pathological changes in many cardiac diseases, however its role in HFpEF was unknown. Here we show that inhibition of *Runx*1 limits adverse cardiac remodelling in a clinically relevant mouse model of HFpEF. Cardiomyocyte-specific tamoxifen-inducible *Runx*1-deficient mice with HFpEF are protected, with preservation of diastolic function, and attenuation of pulmonary oedema, exercise intolerance, and hypertrophy. Furthermore, targeting *Runx*1 in HFpEF by using gene transfer or small molecule inhibitors improves diastolic function, both in female and male mice. Overall, our research enhances our understanding of RUNX1 in cardiac disease and demonstrates a novel translational target for the treatment of HFpEF. Keywords: Heart failure with preserved ejection fraction, metabolic heart failure, diastolic dysfunction, hypertrophy, pulmonary oedema, exercise intolerance

**CLINICAL PERSPECTIVE:** Heart failure (HF) is a leading cause of death world-wide and traditionally divided into different subtypes according to cardiac ejection fraction (EF). In contrast to HF with reduced EF (HFrEF), there are limited treatment options for HF with preserved EF which is of considerable concern given that HFpEF is projected to become the dominant HF subtype in the future ^1^. RUNX1 has been demonstrated to play an important role in the development of many cardiac and non-cardiac diseases. As a result, the potential for RUNX1 inhibitors as therapeutic agents across various conditions has become increasingly evident. In this study we established the therapeutic potential of targeting RUNX1 in the context of HFpEF. Targeting RUNX1 in cardiomyocytes markedly attenuates the development of the HFpEF phenotype and therefore this novel translational therapeutic target has great potential to address one of the biggest challenges in cardiac research.

## INTRODUCTION

Heart failure (HF), a complex syndrome in which the heart is unable to meet the metabolic demands of the body, leads to considerable morbidity and mortality worldwide. It is classically categorised by the proportion of blood ejected from the left ventricle (LV) with each beat, the ejection fraction (EF).

HF with reduced ejection fraction (HFrEF) has been heavily investigated for many years and there are several treatment options available that reduce mortality ^2^. More elusive, however, is HF with *preserved* ejection fraction (HFpEF) which is increasing in prevalence and is poorly understood ^3^.

Despite multiple advances in the treatment of HFrEF, the classic HFrEF treatments are not convincingly effective for use in HFpEF, resulting in limited therapeutic options ^4^. HFpEF is a multimorbidity syndrome, often developing alongside hypertension, metabolic stress, and diabetes, and results in diastolic dysfunction, pulmonary oedema, hypertrophy, and exercise intolerance ^4^.

HFpEF includes a wide range of clinical phenotypes and pathophysiological heterogeneity, and as such it is not clearly understood ^5^. Therefore, elucidating molecular or cellular factors contributing to the development of HFpEF is essential and an important step toward identifying therapeutic targets.

Master-regulator transcription factor, RUNX1, is minimally expressed in the adult heart but can be reactivated in the context of cardiac pathology. Using animal model systems, it has been shown to be a mediator and therapeutic target against adverse cardiac remodelling following myocardial infarction (MI), which is a major cause of HFrEF ^6–8^, and in a transaortic constriction HFrEF model ^9^. Targeting *Runx*1 in the post-MI heart results in improved systolic function, calcium handling, and preservation of genes involved in oxidative phosphorylation ^6,7,10^. Large scale analysis of RNAseq studies on human myocardium ^11–17^ demonstrates that *Runx*1 expression is increased in several cardiac pathologies including myocardial infarction, hypertrophic cardiomyopathy, and dilated cardiomyopathy (Supplemental Figure 1, Supplemental Table 1). Further, serum samples from people admitted to hospital with decompensated HFpEF and HFrEF show that RUNX1 expression is higher in HFpEF than HFrEF (personal communication [Lang/Mooney]). Therefore, we hypothesised a potential role for RUNX1 in the pathophysiology of HFpEF, which is characterised by cardiac hypertrophy and stiffening ^18^. The aim of this study was to use a preclinical model of HFpEF to interrogate the potential role of RUNX1 in the development of HFpEF and to identify its potential as a therapeutic target for the treatment of HFpEF.

## METHODS

Detailed methods and statistical analysis are presented in the supplemental methods. We used a previously established ^19^, two-hit model (2HM) that combines administration of a high-fat diet (HFD) and inhibition of nitric oxide synthase with N^ω^-nitro-L-arginine methyl ester (L-NAME) in drinking water to induce a HFpEF phenotype and compared changes to age-matched controls (CTRL) fed a regular chow diet and normal drinking water ^19^. We utilised cardiomyocyte-specific tamoxifen-inducible *Runx1*-deficient (*Runx1*^Δ/Δ^) mice and floxed genetic-control mice (*Runx1*^fl/fl^) generated as previously described ^20^. RUNX1 was also targeted by either adeno-associated virus (AAV)-mediated delivery of shRNA or a small molecule inhibitor of RUNX1, Ro5-3335, as detailed in the supplemental methods. Bulk RNA sequencing and subsequent pathway analysis was performed on LV tissue samples from *Runx1*^fl/fl^ and *Runx1*^Δ/Δ^ mice at baseline and after the 2HM protocol. Using prior biological knowledge from the Ingenuity knowledge base, the cardiotoxicity networks and functional analyses were generated using QIAGEN Ingenuity Pathway Analysis (IPA) (https://www.qiagenbioinformatics.com/products/ingenuity-pathway-analysis/). The focus on cardiotoxicity processes considers the likely activation and inhibition of biological processes between the two strain comparisons (2HM-*Runx1*^fl/fl^ compared to CTRL-*Runx1*^fl/fl^ and 2HM-*Runx1*^Δ/Δ^ and CTRL-*Runx1*^Δ/Δ^) using a Z-score. Differences in Z-scores were identified.

## RESULTS

### Effect of cardiomyocyte-specific *Runx1-*deficiency on the development of HFpEF

To evaluate the contribution of RUNX1 to the pathophysiology of HFpEF, we utilised male *Runx1*^Δ/Δ^ mice and floxed control mice (*Runx1*^fl/fl^) on the 2HM and CTRL protocols compared to age-matched genetic controls (CTRL-*Runx1*^fl/fl^: n = 19, CTRL-*Runx1*^Δ/Δ^: n = 17, 2HM-*Runx1*^fl/fl^: n = 21, 2HM- *Runx1*^Δ/Δ^: n = 22, Figure 1a). Each of the two-hits were observed in 2HM-*Runx1*^fl/fl^ and 2HM- *Runx1*^Δ/Δ^ male mice with an increase in body weight (8.0 ± 0.9g and 7.0 ± 0.9g, respectively both P<0.05; Figure 1b) and systolic blood pressure (SBP, 33 ± 21mmHg and 30 ± 20mmHg respectively, both P<0.05; Figure 1c) over the course of the protocol compared to respective CTRL groups. We confirmed that neither 2HM group had developed HFrEF by assessing whole heart contractile function as measured by fractional shortening *via* echocardiography (Supplemental Figure 2).

**Figure 1.**
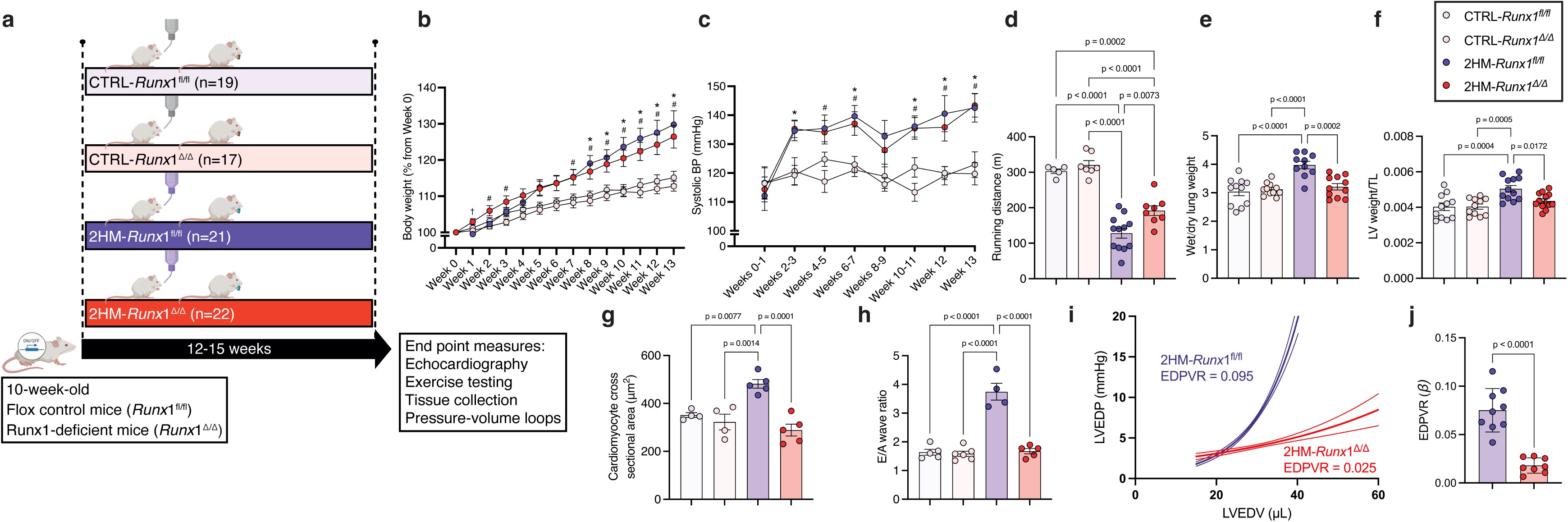
*Runx*1-deficient mice are protected against HFpEF phenotype. **a)** Schematic of two-hit protocol and experimental groups. **b)** Body weight over the experimental protocol in each of the groups. * p < 0.05 for 2HM-*Runx*1^fl/fl^ (n = 21) compared to CTRL-*Runx*1^fl/fl^ (n = 19), # p < 0.05 for 2HM-*Runx*1^Δ/Δ^ (n = 22) compared to CTRL-*Runx*1^Δ/Δ^ (n = 17), and ^†^ p < 0.05 for 2HM-*Runx*1^Δ/Δ^ compared to 2HM-*Runx*1^fl/fl^ by mixed-effects analysis. **c)** Systolic blood pressure (BP) over the experimental protocol. * p < 0.05 for 2HM-*Runx*1^fl/fl^ compared to CTRL-*Runx*1^fl/fl^ and # p < 0.05 for 2HM-*Runx*1^Δ/Δ^ compared to CTRL-*Runx*1^Δ/Δ^ by mixed-effects analysis. Characterisation of the HFpEF phenotype: **d)** Exercise intolerance was quantified by running distance. **e)** Pulmonary oedema quantified by wet to dry lung weight ratio. Hypertrophy was quantified by **f)** left ventricular (LV) weight normalised to tibial length (TL) and by **g)** cardiomyocyte cross sectional area assessed following Wheat Germ Agglutinin staining. Diastolic function quantified by **h)** E to A wave ratio from pulsed wave Doppler echocardiography and by the slope (β) of the end-diastolic pressure volume relationship derived from the exponential equation: (LVEDP= curve fitting constant × e^[stiffness^ ^constant^ ^×^ ^LV^ ^end^ ^diastolic^ ^volume]^) **i)** representative curves (error lines denote 95% confidence interval; and **j)** data set.

We then evaluated other key features of the HFpEF phenotype. At the end of the protocol, 2HM-*Runx1*^fl/fl^ mice had developed exercise intolerance, demonstrated by a reduction in running distance compared to CTRL-*Runx1*^fl/fl^ mice (128 ± 14m *vs*. 296 ± 18m respectively, P<0.05; Figure 1d). 2HM-*Runx1*^Δ/Δ^ mice also had a reduction in running distance compared to their genetic control (2HM-*Runx1*^Δ/Δ^ : 192 ± 14m, CTRL-*Runx1*^Δ/Δ^: 327 ± 22m, P<0.05; Figure 1d) however, the exercise intolerance was attenuated because 2HM-*Runx1*^Δ/Δ^ mice ran greater distances than 2HM-*Runx1*^fl/fl^ mice (128 ± 14m *vs*. 192 ± 14m, P<0.05; Figure 1d). *Runx*1 deficiency also protected mice from developing pulmonary oedema as measured by the wet to dry lung weight. The wet to dry lung weight ratio was increased in 2HM-*Runx1*^fl/fl^ compared to in CTRL-*Runx1*^fl/fl^ mice (3.90 ± 0.2 *vs*. 2.84 ± 0.2, P<0.05; Figure 1e). Conversely, there was no difference between 2HM-*Runx1*^Δ/Δ^ and CTRL-*Runx1*^Δ/Δ^ mice (3.14 ± 0.2 *vs*. 3.00 ± 0.1; Figure 1e), and 2HM-*Runx1*^Δ/Δ^ mice had significantly lower wet to dry lung weight ratio than 2HM-*Runx1*^fl/fl^ mice (3.14 ± 0.2 *vs*. 3.90 ± 0.2, P<0.05; Figure 1e). *Runx*1 deficiency was also protective against development of hypertrophy as measured by LV weight normalised to tibial length (LV/TL). 2HM-*Runx1*^fl/fl^ had increased LV/TL compared to CTRL-*Runx1*^fl/fl^ mice (4.9 ± 2.3*10^-3^ *vs*. 3.5 ± 1.2*10^-3^, P<0.05; Figure 1f) and 2HM-*Runx1*^Δ/Δ^ mice had smaller LV/TL than 2HM-*Runx1*^fl/fl^ mice (4.2 ± 1.3*10^-3^, P<0.05; Figure 1f) but no difference compared to CTRL- *Runx1*^Δ/Δ^ mice (3.7 ± 1.0 *10^-3^; Figure 1f). An additional indicator, relevant to concentric hypertrophy, is cardiomyocyte cross-sectional area. In contrast to 2HM-*Runx1*^fl/fl^ animals, 2HM-*Runx1*^Δ/Δ^ mice showed no significant increase in the cross-sectional area of cardiomyocytes compared to their relative control group (CTRL-*Runx1*^fl/fl^: 352 ± 9.9 μm^2^, CTRL-*Runx1*^Δ/Δ^: 323 ± 32.4 μm^2^, 2HM-*Runx1*^fl/fl^: 482 ± 18.0 μm^2^, 2HM-*Runx1*^Δ/Δ^: 289 ± 24.8 μm^2^; Figure 1g, Supplemental Figure 2). Further, posterior and anterior wall thickness measured during systole with M-mode echocardiography was increased in the 2HM-*Runx1*^fl/fl^ mice but not in the 2HM-*Runx1*^Δ/Δ^ mice, compared to relevant controls (Supplemental Figure 2). In addition to hypertrophy, there was a striking preservation of diastolic function in cardiomyocyte-specific *Runx*1 knockdown mice, quantified by E to A wave ratio from pulsed-wave Doppler echocardiography. Compared to CTRL-*Runx1*^fl/fl^ and CTRL-*Runx1*^Δ/Δ^, 2HM- *Runx1*^fl/fl^ had a higher E/A ratio whereas the E/A ratio in 2HM-*Runx1*^Δ/Δ^ mice was not different from either control group (CTRL-*Runx1*^fl/fl^: 1.63 ± 0.10, CTRL-*Runx1*^Δ/Δ^: 1.58 ± 0.09, 2HM-*Runx1*^fl/fl^: 3.75 ± 0.29, 2HM-*Runx1*^Δ/Δ^: 1.67 ± 0.09; Figure 1h, Supplemental Figure 2). An independent measure of LV chamber stiffness was calculated by fitting the slope of the load-independent end-diastolic pressure- volume relationship (EDPVR) measured using intracardiac pressure-volume catheters. 2HM-*Runx1*^fl/fl^ mice had a steeper EDPVR slope than 2HM-*Runx1*^Δ/Δ^ mice, indicating better diastolic function in the 2HM-*Runx1*^Δ/Δ^ mice (0.075 ± 0.008 *vs*. 0.018 ± 0.003, P<0.05; Figure 1i, 1j). Peripheral organs were also collected to investigate systemic effects of cardiomyocyte-specific *Runx*1-deficiency. Liver, right kidney, and left kidney weights (all normalised to tibial length) were increased in 2HM-*Runx1*^fl/fl^ compared to CTRL-*Runx1*^fl/fl^ mice but were not different between *Runx1*^Δ/Δ^ mouse groups (Supplemental Figure 2).

### *Runx1* RNA interference using adeno-associated virus serotype 9 (AAV9) attenuates diastolic dysfunction in HFpEF

Given the striking phenotypic differences observed in 2HM-*Runx1*^Δ/Δ^ mice compared to 2HM-*Runx1*^fl/fl^, we then tested whether using translational approaches to target RUNX1 expression could prevent the development of HFpEF. To do this we utilised a viral vector-mediated gene delivery approach with AAV9-*Runx*1-shRNA to knockdown *Runx*1 in our 2HM of HFpEF. We injected 12-week-old C57BL/6N male mice via the tail vein with AAV9-scramble-shRNA (2HM-AAV9-scram, n = 10) or AAV9-*Runx*1- shRNA (2HM-AAV9-*Runx*1, n = 11) after which mice were placed on the 2HM protocol for 8 weeks for comparison to age-matched C57BL/6N mice on the 2HM (2HM-C57N, n = 18) or control (CTRL- C57N, n = 13) protocols (Figure 2a). Once again, we confirmed the efficacy of our 2HM by measuring changes in body weight (Figure 2b), SBP (Figure 2c), and preservation of fractional shortening from echocardiograpgy (Supplemental Figure 3) over the duration of the protocol. Targeting *Runx*1 with AAV9-*Runx*1-shRNA was effective in preventing a number (but not all) of the key features of the HFpEF phenotype. Exercise intolerance was observed in all three 2HM groups compared to CTRL- C57N, but with no difference in running distance between 2HM groups (CTRL-C57N: 262 ± 12m, 2HM-C57N: 108 ± 3m, 2HM-AAV9-scram: 108 ± 23m, 2HM-AAV9-*Runx*1: 163 ± 22m; Figure 2d).

**Figure 2.**
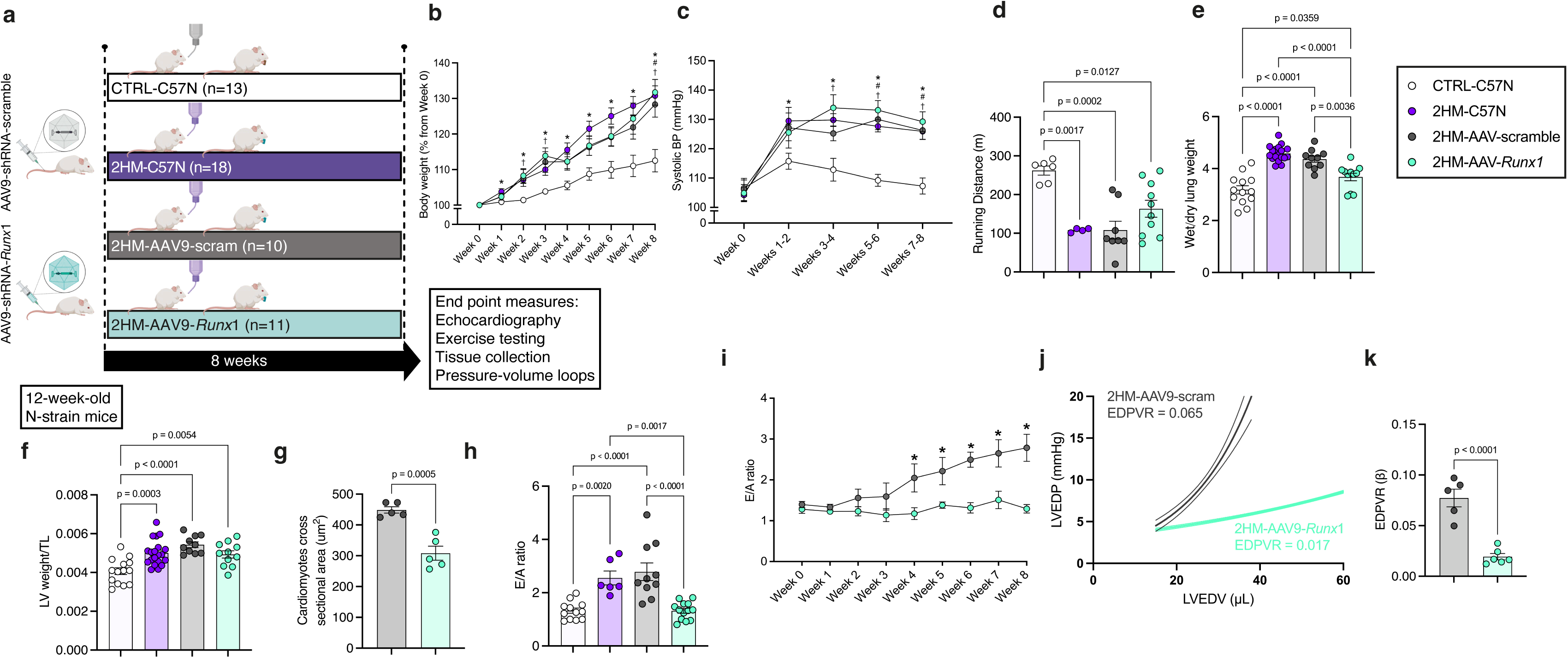
AAV9-mediated knockdown of *Runx*1 protects against diastolic dysfunction. **a)** Schematic of two-hit protocol and experimental groups. **b)** Body weight over the experimental protocol i* p < 0.05 for 2HM-C57N (n = 18) compared to CTRL-C57N (n = 13), # p < 0.05 for 2HM-AAV9-scraM (n = 10) compared to CTRL-C57N, and ^†^ p < 0.05 for 2HM-AAV9-*Runx*1 (n = 11) compared to CTRL-C57N by mixed-effects analysis. **c)** Systolic blood pressure (BP) over the experimental protocol. * p < 0.05 for 2HM-C57N compared to CTRL-C57N, # p < 0.05 for 2HM-AAV9-scram compared to CTRL-C57N, and ^†^ p < 0.05 for 2HM-AAV9-*Runx*1 compared to CTRL-C57N by mixed-effects analysis. Characterisation of the HFpEF phenotype: **d)** Exercise intolerance was quantified by running distance. **e)** Pulmonary oedema quantified by wet to dry lung weight ratio. Hypertrophy was quantified by **f)** left ventricular (LV) weight normalised to tibial length (TL) and by **g)** cardiomyocyte cross sectional area assessed following Wheat Germ Agglutinin staining. Diastolic function quantified by E to A wave ratio from pulsed wave Doppler echocardiography on **h)** the final week and **i)** over the protocol; and by the slope (β) of the end-diastolic pressure volume relationship derived from the exponential equation: (LVEDP= curve fitting constant × e^[stiffness^ ^constant^ ^×^ ^LV^ ^end^ ^diastolic^ ^volume]^) **j)** representative curves (error lines denote 95% confidence interval; and **k)** data set.

AAV9-*Runx*1 did, however, attenuate the development of pulmonary oedema compared to the other two 2HM groups, quantified by wet to dry lung weight ratio (CTRL-C57N: 3.18 ± 0.16, 2HM-C57N: 4.58 ± 0.07, 2HM-AAV9-scram: 4.36 ± 0.12, 2HM-AAV9-*Runx*1: 3.68 ± 0.15; Figure 2e). As with exercise testing, LV/TL was increased in 2HM groups compared to CTRL but was not different between 2HM groups (CTRL-C57N: 4.1 ± 0.2 *10^-3^, 2HM-C57N: 5.0 ± 0.2 *10^-3^, 2HM-AAV9-scram: 5.4 ± 0.2 *10^-3^, 2HM-AAV9-*Runx*1: 4.9 ± 0.2 *10^-3^; Figure 2f). However, cardiomyocyte cross-sectional area of 2HM-AAV9-Runx1 was less than the 2HM-AAV9-scram group (2HM-AAV9-scram: 449 ± 10 μm^2^ *vs*. 2HM-AAV9-*Runx*1: 308 ± 23 μm^2^; Figure 2g, Supplemental Figure 3). Most striking was the preservation of diastolic function by targeting *Runx*1 with AAV9. E/A ratio was increased in both 2HM- C57N and 2HM-AAV9-scram groups compared to CTRL-C57N but was not increased in 2HM-AAV9- *Runx*1 compared to CTRL-C57N (CTRL-C57N: 1.32 ± 0.10, 2HM-C57N: 2.55 ± 0.26, 2HM-AAV9-scram: 2.78 ± 0.33, 2HM-AAV9-*Runx*1: 1.29 ± 0.12; Figure 2h, 2i; the latter figure demonstrating change over time, Supplemental Figure 3). This was also consistent with EDPVR, which was markedly lower in the 2HM-AAV9-*Runx*1 group compared to 2HM-AAV9-scram (0.077 ± 0.009 *vs*. 0.019 ± 0.003, P<0.05; Figure 2j, 2i).

### Small molecule inhibition of RUNX1 remedies the HFpEF phenotype

To take this one translational step further, we aimed to identify if inhibition of RUNX1 could ameliorate the HFpEF phenotype once it has already begun to develop using an established small molecule inhibitor of RUNX1 ^20^. 10-12 week-old C57BL/6N strain male mice were placed on the 2HM protocol for 10-12 weeks. Prior to drug treatment, *in vivo* parameters were utilised to ensure the HFpEF phenotype had developed and any mice that did not have HFpEF symptoms were excluded so that we were only attempting to treat mice with a phenotype to attenuate. Next, while mice remained on 2HM protocol, we injected small molecule inhibitors of RUNX1, either DMSO or Ro5-3335 every second day for two weeks prior to collecting end-point measurements and organometrics (Figure 3a). Although RUNX1 inhibition by Ro5-3335 injections did not change exercise tolerance (199.1 ± 50.2 *vs*. 192.5 ± 40.21, P>0.05; Figure 3b), pulmonary oedema was reduced in 2HM-Ro5-3335 mice compared to 2HM-DMSO mice (4.26 ± 0.05 *vs*. 4.06 ± 0.07, P<0.05; Figure 3c). As with exercise intolerance, hypertrophy was not changed by Ro5-3335 administration (4.9x10^-3^ ± 1.1x10^-4^ *vs*. 5.07x10^-3^ ± 1.476x10^-4^, P>0.05, Figure 3d). Diastolic dysfunction was attenuated in 2HM-Ro5-3335 mice compared to 2HM-DMSO. There was no difference in E/A wave ratio post-injection compared to pre-injection in the 2HM-DMSO mice (2.66 ± 0.35 *vs*. 2.82 ± 0.15, P>0.05; Figure 3e, left, Supplemental Figure 4) whereas the post-injection E/A wave ratio was reduced in the 2HM-Ro5-3335 mice compared to pre-injection, demonstrating diastolic dysfunction was attenuated (1.68 ± 0.14 *vs*. 2.51 ± 0.16, P<0.05; Figure 3e, right, Supplemental Figure 4). This was confirmed using PV loop assessment of diastolic function by EDPVR in 2HM-DMSO mice compared to the 2HM-Ro5-3335 group (0.048 ± 0.005 *vs*. 0.029 ± 0.005, P<0.05; Figure 3f).

**Figure 3.**
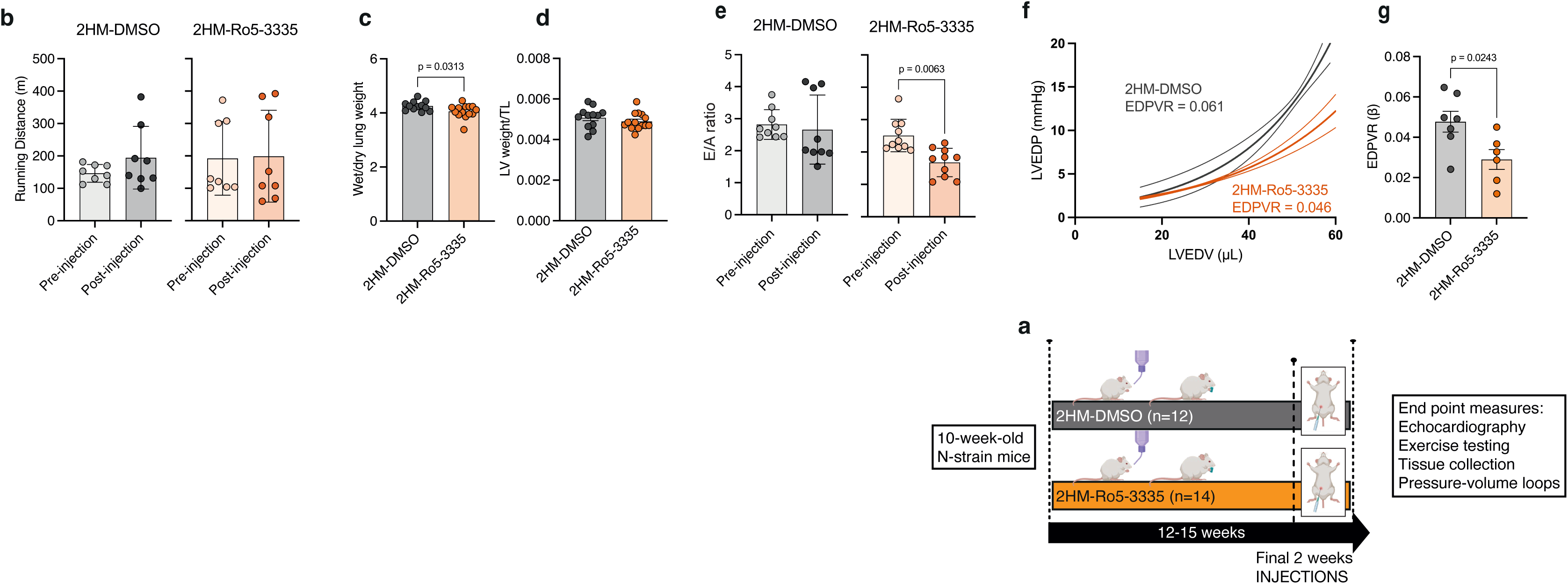
*Runx*1 small molecule inhibitor Ro5-3335 remedies against diastolic dysfunction. **a)** Schematic of two-hit protocol and experimental groups, 2HM-DMSO (n = 12) and 2HM-Ro5-3335 (n = 14). Characterisation of the HFpEF phenotype: **b)** Exercise intolerance was quantified by running distance. **c)** Pulmonary oedema was quantified by wet to dry lung weight ratio. Hypertrophy was quantified by **d)** left ventricular (LV) weight normalised to tibial length (TL). Diastolic function quantified by **e)** E to A wave ratio from pulsed wave Doppler echocardiography and by the slope (β) of the end-diastolic pressure volume relationship derived from the exponential equation: (LVEDP= curve fitting constant × e^[stiff-ness^ ^constant^ ^×^ ^LV^ ^end^ ^diastolic^ ^volume]^) **f)** representative curves (error lines denote 95% confidence interval; and **g)** data set.

### RNAseq predicts patterns of transcriptional changes consistent with a HFpEF phenotype

To gain broader insight into the role of *Runx*1 in HFpEF, we performed bulk RNAseq on analysis on LV tissue samples from *Runx1*^fl/fl^ and *Runx1*^Δ/Δ^ mice both at baseline (day 0, D0) and at the end of the 2HM study. There were not any significantly differentially expressed genes (DEG) between *Runx1*^fl/fl^ and *Runx1*^Δ/Δ^ at D0 and despite the large phenotypic differences, there were only 32 DEG between *Runx1*^fl/fl^ and *Runx1*^Δ/Δ^ mice at week 13 (Supplemental Table 2). However, there were many differences when comparing each strain at week 13 compared to their respective baseline controls. Thus, because the transcriptomic snapshot at the end of the study does not depict the highly different phenotypes, we focussed on comparing the changes from D0 to the end time point within each strain. Using a false discovery rate (FDR) cut-off of ≤0.05 and log fold change (logFC) ±1, there were 1,866 DEG in 2HM-*Runx*1*^fl/fl^* mice at week 13 compared to D0 *Runx*1*^fl/fl^* mice (Figure 4a and 4c) and 3,691 DEG at week 13 in 2HM-*Runx*1^Δ/Δ^ mice compared to the D0 (Figure 4b and 4c). The majority of DEG were shared between strains (1727 DEG: 92.6% of total DEG for *Runx*1*^fl/fl^* and 53.2% of total DEG for *Runx*1^Δ/Δ^; Figure 4c). Interestingly, the unique changes in the *Runx*1^Δ/Δ^ mice across timepoints may account for the large functional differences observed because there were very few unique changes in the Runx1*^fl/fl^* mice (Figure 4c). Using all significantly DEG in *Runx1*^fl/fl^ and *Runx1*^Δ/Δ^ mice at week 13 compared to baseline, we focused on cardiac toxicity functions defined by IPA software. We compared Z-scores (a statistical measure utilised to determine the significance and directionality of gene expression changes within a given pathway over that time course) from *Runx1*^fl/fl^ (week 13 of 2HM *vs* D0) and *Runx1*^Δ/Δ^ (week 13 of 2HM *vs* D0). We visualised the impact of *Runx*1 deficiency by plotting the difference between Z-scores from Runx1*^fl/fl^* (2HM *vs* D0) minus Runx1^Δ/Δ^ (2HM *vs* D0) mice (Z-diff; Figure 4d). Whilst some predictive changes in cardiac toxicity functions demonstrated limited difference in Z-diff (yellow; Figure 4d), the largest differences in Z-score were in congestive heart failure genes and cardiac damage genes. The genes included by IPA in these two cardiac toxicity functions were then plotted using a heat map (Figure 4e and f).

**Figure 4.**
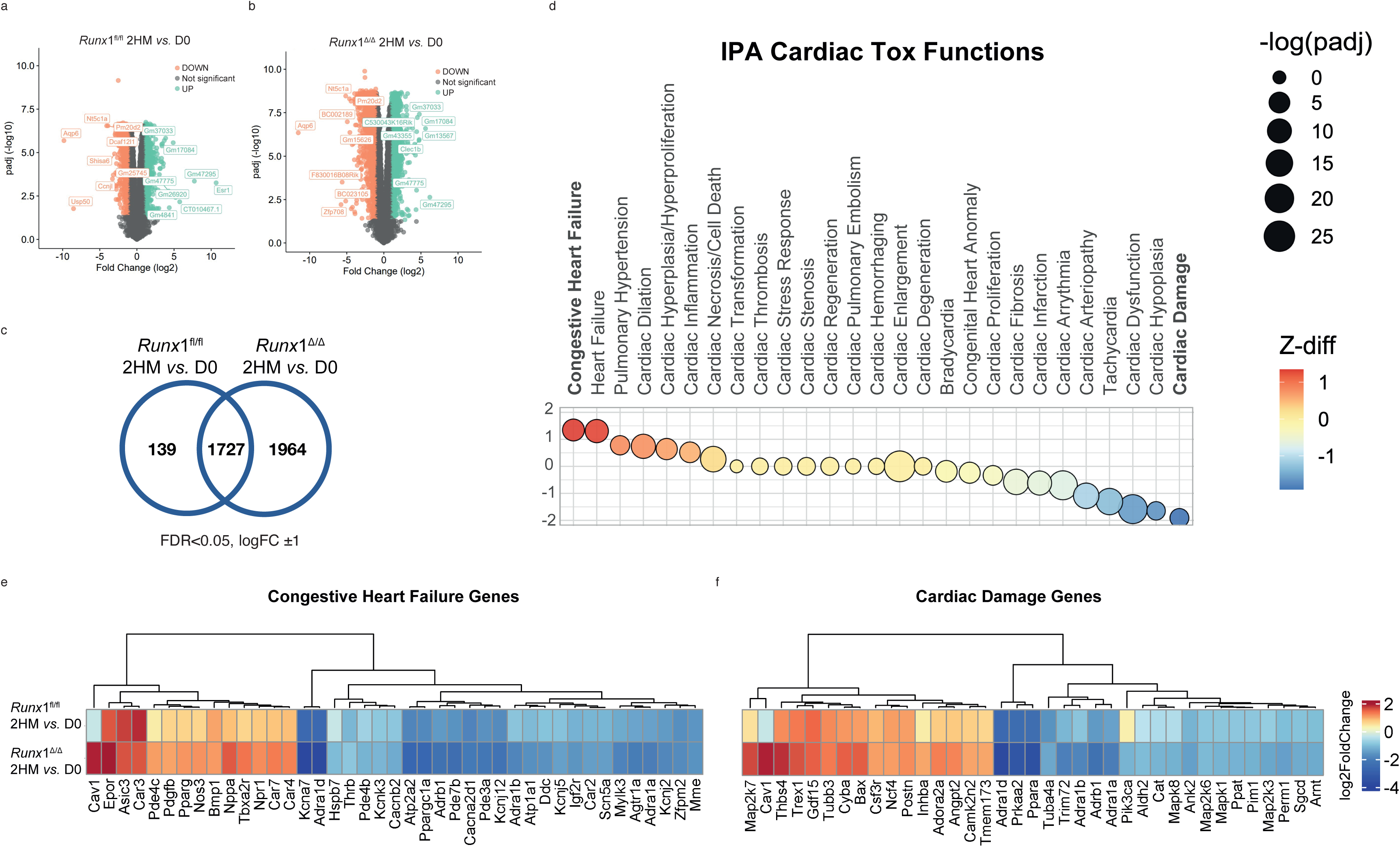
Cardiac differential gene expression (DEG) analysis of *Runx*1^fl/fl^ and *Runx*1^Δ/Δ^ mice between day 0 (D0) and 13 weeks of two-hit model (2HM) protocol. Volcano plots of all genes (orange-significantly downregulated, green-significantly upregulated and grey not changing) with the eight most regulated genes indicated in **a)** *Runx*1^fl/fl^ and **b)** *Runx*1^Δ/Δ^ mice. **c)** Venn diagram indicating unique changes and the large number of genes that are commonly differentially regulated between group comparison. **d)** Differences in functional predictions using Z-scores comparing *Runx*1^fl/fl^ mice to *Runx*1^Δ/Δ^ mice (red indicating an activation between *Runx*1^fl/fl^ minus *Runx*1^Δ/Δ^, blue indicating an inhibition, and yellow indicating similar functional predictions in both groups). Heat map representing patterns of **e)** congestive heart failure and **f)** cardiac damage gene expression levels between *Runx*1^fl/fl^ mice to *Runx*1^Δ/Δ^ mice.

### Inhibition of *Runx*1 in female mice: reversal of HFpEF phenotype

To further increase the relevance and impact of our findings, we expanded our study in two ways: we used female mice to increase clinical relevance; and we waited to intervene with *Runx*1 inhibition *via* RNA interference until HFpEF was already established in the mice, to test its utility as a therapy. It has been demonstrated that it is more difficult to induce a HFpEF phenotype *via* the 2HM in female mice compared to males in young mice ^21^. Thus, in a cohort of C57-N strain females we waited until they were aged 14 weeks (∼40% older than previous data) before placing them on the 2HM protocol with a ramping dose of L-NAME (Figure 5a). Once again, we ensured efficacy of the two hits by measuring body weight and SBP in a female CTRL-C57N group (F-CTRL-C57N, n = 4) compared to a female 2HM-C57N group (F-2HM-C57N, n = 16; Figure 5b, 5c). We utilised our intermediary *in vivo* phenotypic measures exercise intolerance (Figure 5d) and diastolic dysfunction (Figure 5e) to confirm that at the 8-week time point the F-2HM-C57N group had established a HFpEF phenotype. Following this, we split the F-2HM-C57N group into two groups for AAV-mediated gene delivery such that they had consistent starting parameters (Figure 5a). One group was injected with AAV9-scramble-shRNA (F-2HM-AAV9-scram, n = 8) and a second injected with AAV9-*Runx*1-shRNA to knockdown *Runx*1 (F-2HM-AAV9-*Runx*1, n = 8). Consistent with the male AAV study, there were no differences in running distance between groups 4 weeks following AAV injection (F-2HM-AAV9-scram: 188 ± 8m, F-2HM-AAV9-*Runx*1: 182 ± 17m, p = 0.7560; Figure 5f). We found pulmonary oedema was reduced in the F-2HM-AAV9-*Runx*1 compared to F-2HM-AAV9-scram (3.97 ± 0.05 *vs* 4.29 ± 0.07, respectively, p = 0.0024; Figure 5h) which was consistent with the male data (Figure 2e). Although hypertrophy (measure by LV weight normalised to TL) was not different between the 2HM-AAV-scram and 2HM-AAV-*Runx*1 males (Figure 2f), it was reduced in F-2HM-AAV9-*Runx*1 compared to F-2HM-AAV-scram (3.1 ± 0.1 *10^-3^ *vs* 3.6 ± 0.2 *10^-3^, respectively, P<0.05; Figure 5i). Finally, diastolic dysfunction was attenuated as measured both by E/A wave ratio from pulse wave Doppler echocardiography (F-2HM- AAV-scram: 1.99 ± 0.11 *vs* F-2HM-AAV9-*Runx*1: 1.49 ± 0.05, p = 0.0007 Figure 5j, Supplemental Figure 5), and by the slope of the EDPVR (F-2HM-AAV-scram: 0.082 ± 0.0002 *vs* F-2HM-AAV9- *Runx*1: 0.041 ± 0.0057, p = 0.0027; Figure 5k).

**Figure 5.**
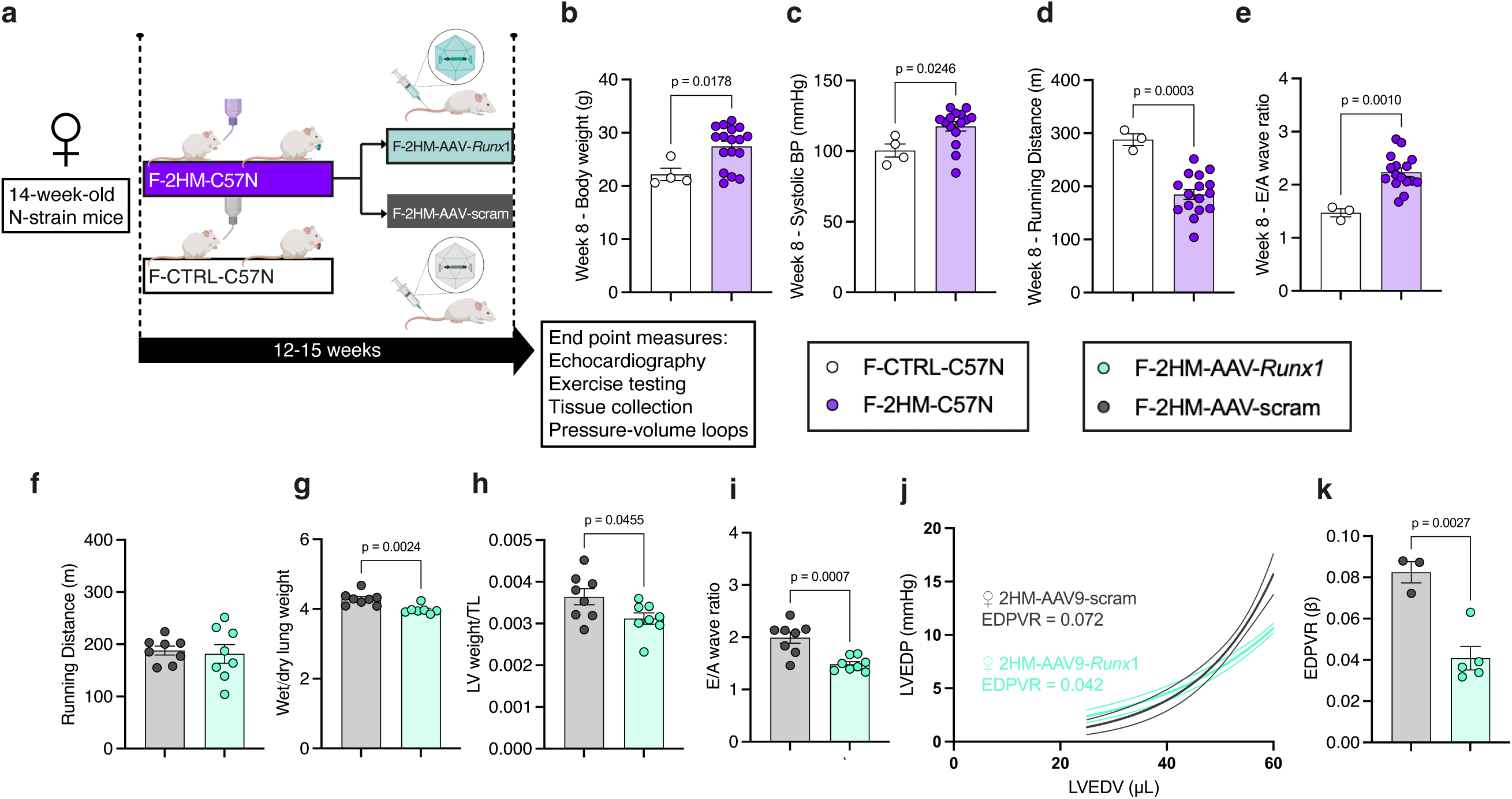
AAV9-mediated knockdown of *Runx*1 in female mice: partial reversal of HFpEF phenotype. **a)** Schematic of two-hit protocol and experimental groups, F-CTRL-C57N (n = 4), F-2HM-C57N (n = 16: F-2HM-AAV-scram, n = 8; F-2HM-AAV-*Runx*1, n = 8). **b)** Body weight, **c)** systolic blood pressure (BP), **d)** exercise intolerance testing quantified by running distance, and diastolic function quantified by **e)** E to A wave ratio from pulsed wave Doppler echocardiography at week 8 in female control C57N-strain mice (F-CTRL-C57N) compared to female C57N-strain mice on the two hit model protocol (F-2HM-C57N). **f)** Exercise intolerance was quantified by running distance and **g)** pulmonary oedema was quantified by **h)** wet to dry lung weight ratio following removal of outlier identified by ROUT outlier test. Diastolic function quantified by **i)** E to A wave ratio from pulsed wave Doppler echocardiography; and by the slope (β) of the end-diastolic pressure volume relationship derived from the exponential equation: (LVEDP= curve fitting con-

## DISCUSSION

This work identifies a critical role for RUNX1 in the development of HFpEF. Furthermore, we provide evidence that targeting *Runx*1 in the context of HFpEF has clinical translational potential.

In recent years, significant work has been done to establish a model with preserved EF which not only demonstrates increased hypertrophy but also phenotypes such as pulmonary oedema, exercise intolerance, and diastolic dysfunction and therefore is more representative of the multimorbidity, multi-system disorder of HFpEF in humans ^19,22^.

*Runx*1 has been robustly demonstrated to play an important role in the context of cardiac disease, with a particular emphasis on its importance in adverse cardiac remodelling following MI ^10,20,23,24^. Previous work has demonstrated the beneficial effects of targeting *Runx*1 in the context of acute MI ^20^ and in the context of ischemic heart disease, however whether these benefits would be observed in a cardiac disease of a chronic progressive nature such as HFpEF was unknown.

Therefore, we adapted a 2HM of HFpEF in our line of transgenic mice with cardiomyocyte *Runx*1 deficiency, and then again with the C57-N strain mice using translational approaches to target *Runx*1. Overall, this work has identified RUNX1 as a promising therapeutic target for treatment and prevention of HFpEF.

Targeting *Runx*1 with a cardiomyocyte-specific *Runx*1-deficient mouse attenuates the development of a HFpEF phenotype. *Runx*1-deficiency is highly protective against the development of HFpEF because despite the efficacy of the two-hits (*i.e.*, mice in both 2HM groups gained weight and had increased SBP), the *Runx*1-deficient mice did not develop all the signs of HFpEF whereas control mice had a classical HFpEF phenotype. Specifically, *Runx*1-deficiency reduced the development of hypertrophy and exercise intolerance and completely protected against development of pulmonary oedema and diastolic dysfunction. Although in this study we have simply targeted a single gene (*Runx*1) in a single cell type (cardiomyocytes), the phenotypic outcome was evident systemically including effects on exercise intolerance, pulmonary oedema, and the mass of peripheral organs, reflecting the beneficial effects of targeting *Runx*1 for both cardiac dysfunction and peripheral systems.

We corroborated and translated these findings using RNAi therapy and small molecule inhibition of RUNX1. Interestingly, similar to the convincing protection of *Runx*1-deficient mice, targeting *Runx*1 with RNAi and small molecule inhibitors also attenuated diastolic dysfunction and pulmonary oedema. Not only were we able to prevent these two phenotypes with pre-treatment of AAV9 targeting *Runx*1 but also by inhibiting *Runx* with the small molecule inhibitor Ro5-3335 after the establishment of HFpEF. We note that some parameters measured were less affected by these alternative approaches and may reflect the number of cardiomyocytes exposed to the therapy and/or, effects on non-cardiomyocytes or duration of exposure. Future work will aim to further understand the relative benefits of different approaches.

To interrogate potential gene changes underlying the phenotypic differences observed when inhibiting *Runx*1, we used RNAseq. This resulted in predictions using IPA software for changes in the regulation of diseases and functions when comparing the final time point (after 13 weeks of 2HM) tissue in both groups compared to day 0 (D0) heart tissue. In any chronic disease it is difficult to determine at which timepoint transcriptional changes might best be identified in order to discern differences between *Runx1*^fl/fl^ and *Runx1*^Δ/Δ^ mice because relevant changes in the transcriptome may precede phenotype differences. As such, it is perhaps unsurprising that the most DEG were shared between strains despite the stark phenotypic differences between the 2HM transgenic groups. However, IPA did predict an upregulation in congestive heart failure pathways in both *Runx1*^fl/fl^ and *Runx1*^Δ/Δ^ mice, with larger changes occurring in *Runx1*^fl/fl^ compared to *Runx1*^Δ/Δ^ mice despite more gene changes overall occurring in the *Runx1*^Δ/Δ^ mice. Interestingly, it was predicted that the upregulation of cardiac damage pathways would result in larger changes in *Runx1*^Δ/Δ^ mice compared to *Runx1*^fl/fl^ mice.

Finally, we expanded the translational relevance of our work by performing a study in female mice. This enabled us to not only determine the effect of gene transfer in both sexes but also interrogate the translational potential of targeting *Runx*1 with AAV9 <u>after</u> the HFpEF phenotype was fully established (in contrast to our male study where AAV was administered prior to mice being placed on the 2HM protocol). The AAV9 was injected following development of an evident HFpEF phenotype. Overall, inhibiting *Runx*1 via RNAi was effective in reducing hypertrophy, pulmonary oedema, and diastolic dysfunction in the female 2HM, thus indicating a potential role for *Runx*1 in the treatment of HFpEF in both females as well as males, and is capable of partially reversing the phenotype.

Limitations to this study include the use of bulk RNAseq rather than a more targeted approach. It is possible that many of the DEG in our bulk tissue samples will be the result of transcriptional changes in non-cardiomyocyte cell types in the ventricle, potentially diluting cardiomyocyte-specific changes that are the result of the *Runx*1-deficiency. Single-cell transcriptomic analysis, potentially at multiple time points, is part of the programme of future work.

The aetiology of HFpEF and the associated changes in heart structure and diastolic function are complex and relatively poorly understood. The relative contributions of the metabolic changes at a cellular level and the chronic low-grade inflammation that accompanies metabolic stress and hypertension are not clear. It is remarkable that the relatively simple model developed by Schiattarella *et al* in 2019 and used again here can recapitulate many of the phenotypic changes associated with HFpEF given the subtleties of these physiological insults and their complex interplay. We acknowledge that RUNX1 is likely to modulate several aspects that mediate the pathogenesis of this syndrome. Interestingly, this is not at the level of the inducing factors, because weight gain and increased SBP are observed in both the *Runx1*^fl/fl^ and *Runx1*^Δ/Δ^ groups. Rather, it appears that RUNX1 is involved in the pathways that connect these factors (metabolic stress and increased SBP) to changes in the heart, leading to hypertrophy and diastolic dysfunction. It is intriguing that attenuation of *Runx*1 function alleviates the deleterious effects observed both following MI and prevents and reverses key aspects of HFpEF, hinting at a more fundamental role in the response of heart tissue to damage and pathophysiological insult.

Overall, this study clearly demonstrates that RUNX1 drives pathological changes in cardiomyocytes in the context of HFpEF. Inhibition of *Runx*1 by gene transfer or the use of a small molecule inhibitor improves LV diastolic function and represents an exciting translational approach for the treatment of HFpEF.

## ACKNOWLEDGMENTS

The authors thank Michael Dunne, Margaret Bell, and Catherine Hawksby and the Biological Services staff from the University of Glasgow Cardiovascular Research Unit for their surgical, animal, and technical assistance. We thank Douglas Strathdee, the Transgenic Technologies Lab, BSU and Histology lab at the Beatson Institute.

## FUNDING

This work was supported by a BHF programme grant RG/20/6/35095 to C.L., E.C, S.N. and C. B. C.B. and C.M.L were also supported by the British Heart Foundation (RE/18/6/34217).

## DISCLOSURE OF INTEREST

Authors have nothing to disclose.

## DATA AVAILABILITY STATEMENT

Data from this study are available upon request to the corresponding author.

---

**SUPPLEMENTAL FIGURE 1.**
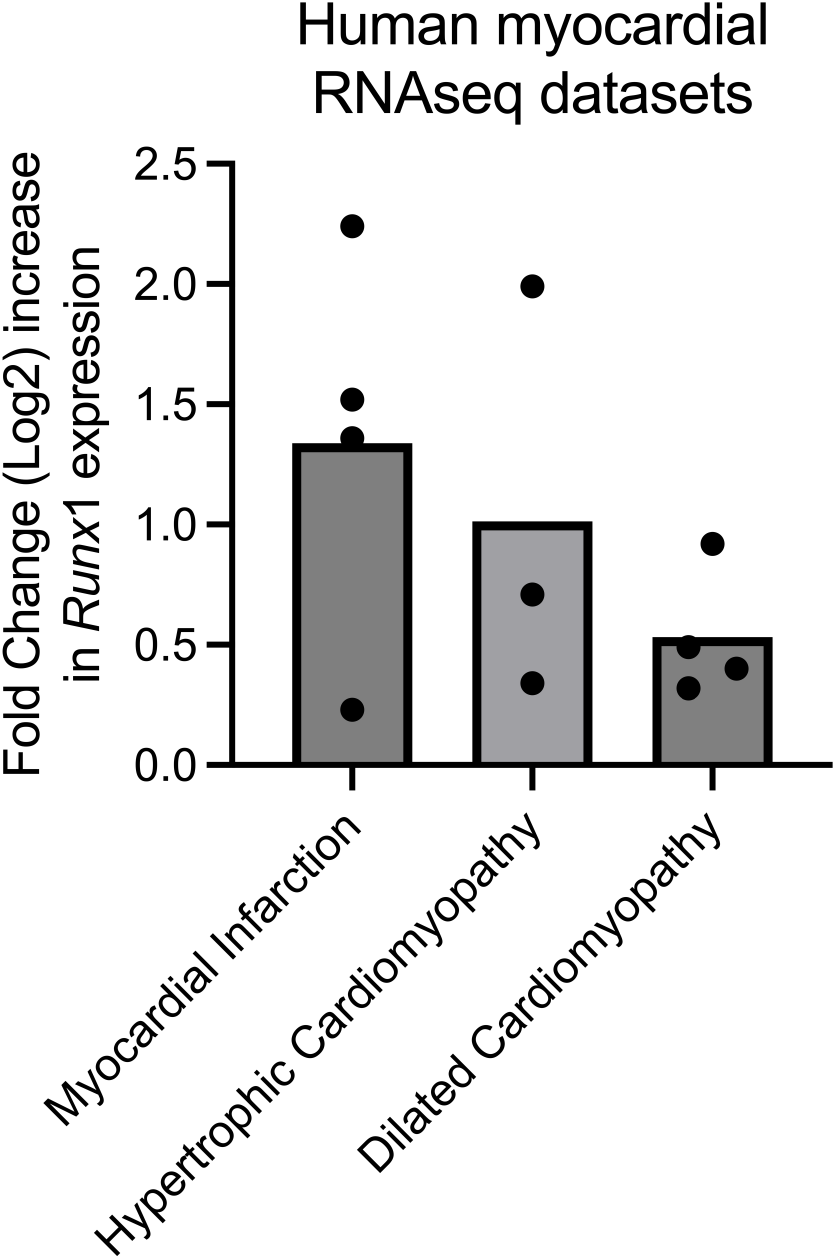

**SUPPLEMENTAL FIGURE 2.**
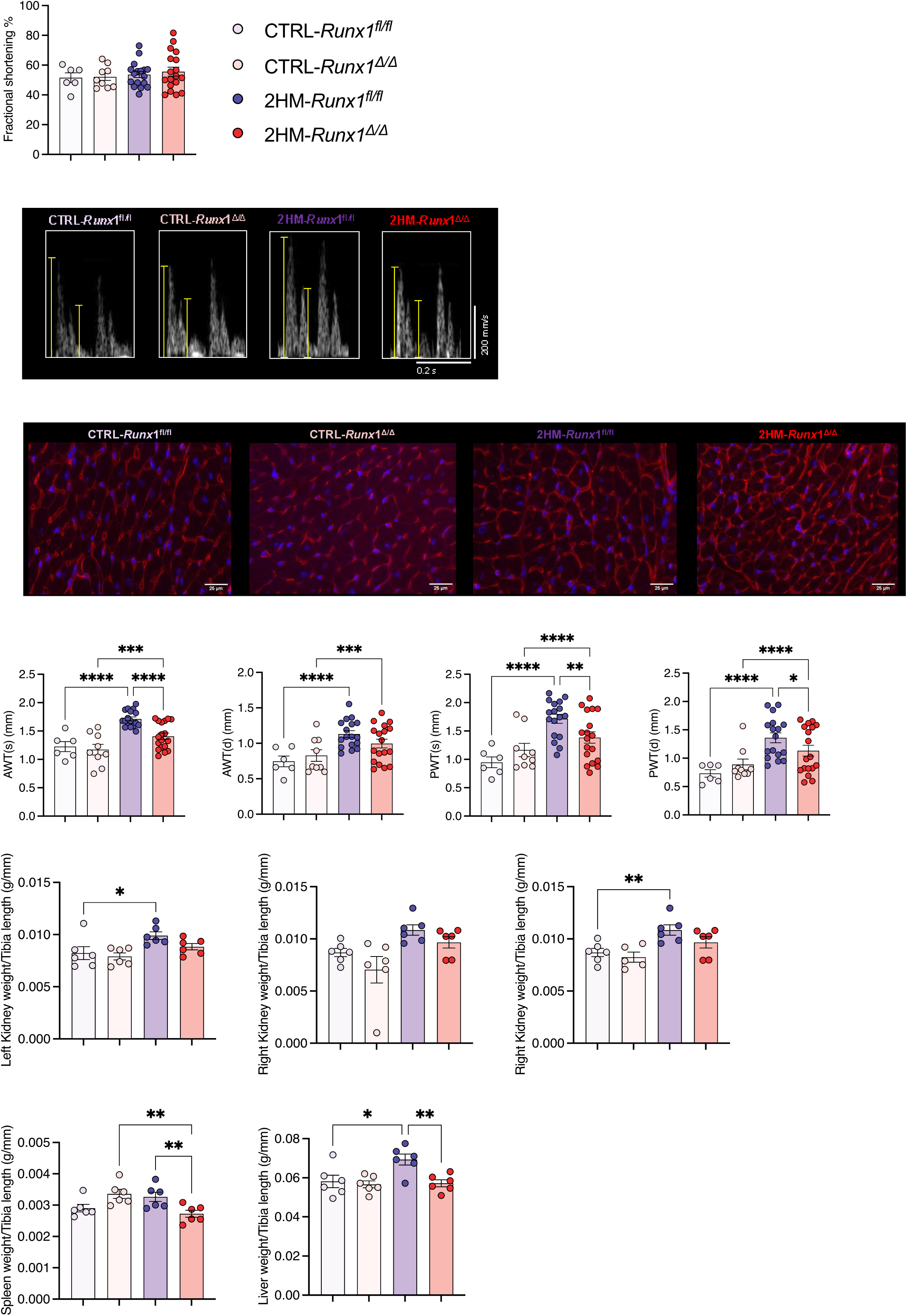

**SUPPLEMENTAL FIGURE 3.**
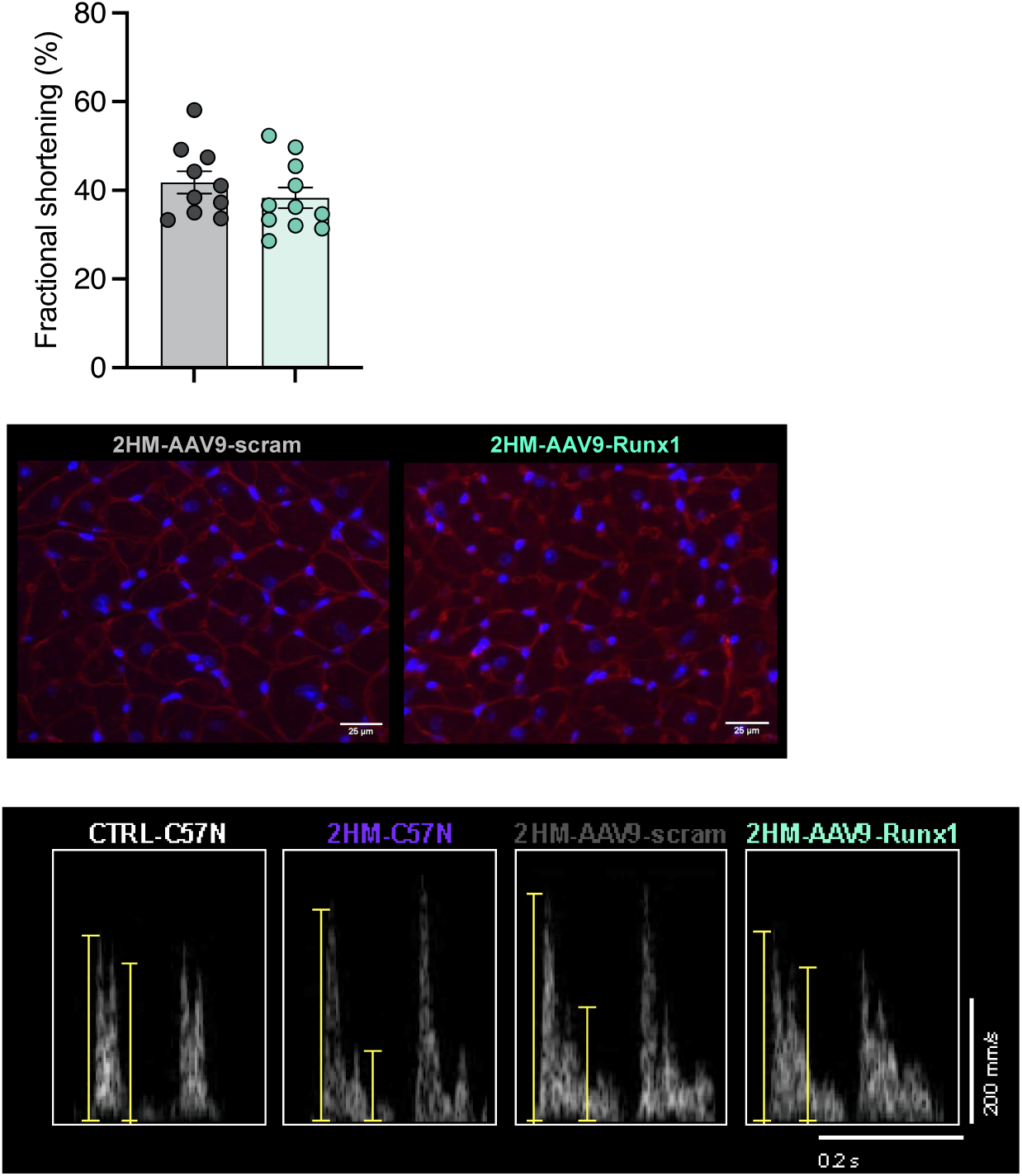

**SUPPLEMENTAL FIGURE 4.**
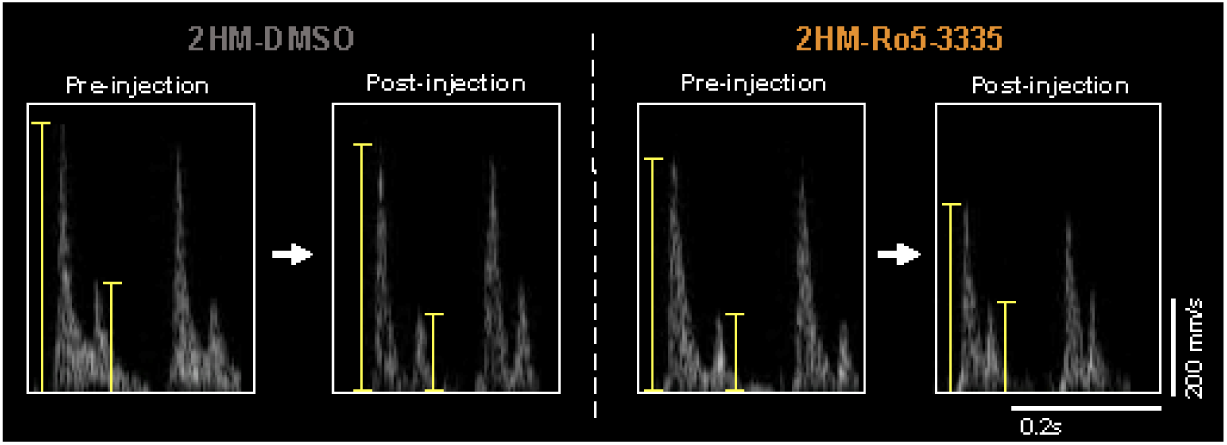

**SUPPLEMENTAL FIGURE 5.**
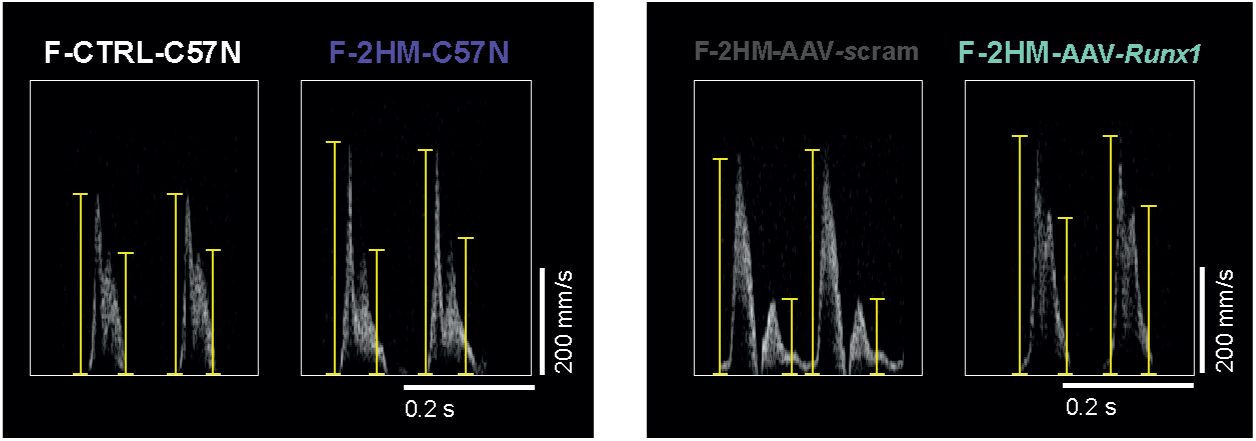

**SUPPLEMENTAL TABLE 1.**

| ComparisonID | GeneName | ComparisonContrast | Pvalue | Adjusted Pvalue | Fold Change [Log2] | Fold change (UP) | Case.DiseaseState |
| --- | --- | --- | --- | --- | --- | --- | --- |
| Myocardial Infarction |  |  |  |  |  |  |  |
| GSE120852.GPL16791.DESeq2R.test3 | RUNX1 | ExperimentGroup => right ventricular tissue from ICM, biventricular heart failure vs right v | 0.0005 | 0.0097 | 2.24 | 4.74 | heart failure;myocardial ischemia |
| GSE116250.GPL16791.DESeq2R.test7 | RUNX1 | Gender:DiseaseState => male -> myocardial ischemia vs disease control | 0.0019 | 0.011 | 1.52 | 2.88 | myocardial ischemia |
| GSE116250.GPL16791.DESeq2R.test2 | RUNX1 | DiseaseState => myocardial ischemia vs disease control | 0.0014 | 0.0072 | 1.36 | 2.57 | myocardial ischemia |
| GSE1145.GPL570.test4 | RUNX1 | DiseaseState => myocardial ischemia vs normal control | 0.0009 | 0.0063 | 0.23 | 1.18 | myocardial ischemia |
| Hypertrophic Cardiomyopathy |  |  |  |  |  |  |  |
| GSE89714.GPL11154.DESeq2R.test1 | RUNX1 | DiseaseState => hypertrophic cardiomyopathy vs normal control | 1.41E-08 | 8.01E-07 | 1.99 | 3.98 | hypertrophic cardiomyopathy |
| GSE141910.GPL16791.DESeq2R.test2 | RUNX1 | DiseaseGroup => hypertrophic cardiomyopathy vs non-failing donor | 0.0006 | 0.0033 | 0.71 | 1.64 | hypertrophic cardiomyopathy |
| GSE1145.GPL570.test2 | RUNX1 | DiseaseState => hypertrophic cardiomyopathy vs normal control | 0.0016 | 0.0152 | 0.34 | 1.27 | hypertrophic cardiomyopathy |
| Dilated Cardiomyopathy |  |  |  |  |  |  |  |
| GSE35108.GPL6244.test4 | RUNX1 | Transfection:DiseaseState => GFP -> dilated cardiomyopathy vs normal control | 0.0002 | 0.0021 | 0.92 | 1.89 | dilated cardiomyopathy |
| GSE35108.GPL6244.test3 | RUNX1 | Transfection:DiseaseState => ATP2A2 -> dilated cardiomyopathy vs normal control | 0.0022 | 0.0095 | 0.49 | 1.41 | dilated cardiomyopathy |
| GSE141910.GPL16791.DESeq2R.test1 | RUNX1 | DiseaseGroup => dilated cardiomyopathy;heart failure vs non-failing donor | 0.0004 | 0.0015 | 0.40 | 1.32 | dilated cardiomyopathy;heart failure |
| GSE155495.GPL21697.DESeq2R.test1 | RUNX1 | DiseaseState => dilated cardiomyopathy;heart failure vs disease control | 0.0121 | 0.0436 | 0.32 | 1.25 | dilated cardiomyopathy;heart failure |

**SUPPLEMENTAL TABLE 2.**
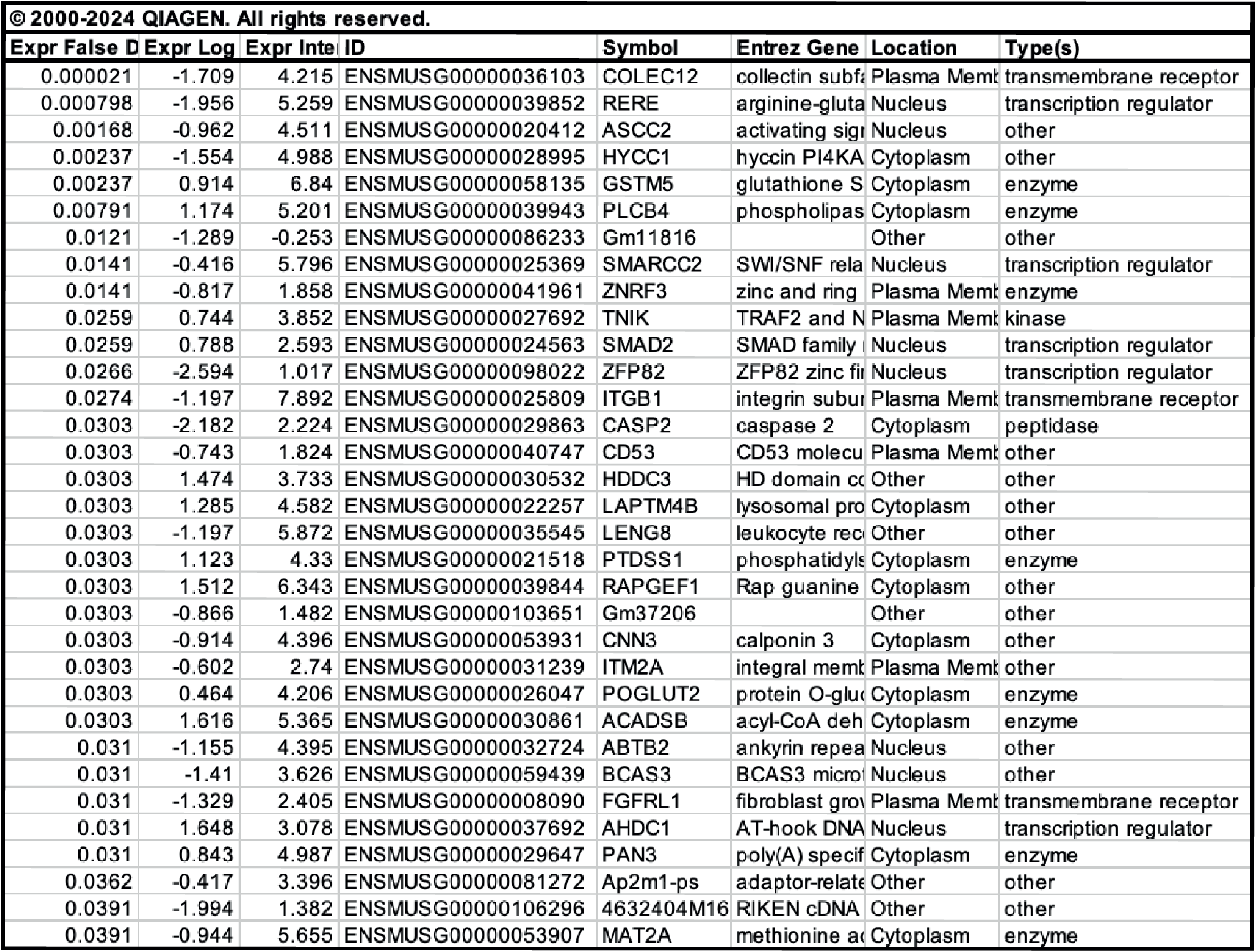

| © 2000-2024 QIAGEN. All rights reserved. |  |  |  |  |  |  |  |
| --- | --- | --- | --- | --- | --- | --- | --- |
| Expr False D | Expr Log | Expr Inte | ID | Symbol | Entrez Gene | Location | Type(s) |
| 0.000021 | -1.709 | 4.215 | ENSMUSG00000036103 | COLEC12 | collectin subfa | Plasma Memk | transmembrane receptor |
| 0.000798 | -1.956 | 5.259 | ENSMUSG00000039852 | RERE | arginine-gluta | Nucleus | transcription regulator |
| 0.00168 | -0.962 | 4.511 | ENSMUSG00000020412 | ASCC2 | activating sig | Nucleus | other |
| 0.00237 | -1.554 | 4.988 | ENSMUSG00000028995 | HYCC1 | hyccin PI4KA | Cytoplasm | other |
| 0.00237 | 0.914 | 6.84 | ENSMUSG00000058135 | GSTM5 | glutathione S | Cytoplasm | enzyme |
| 0.00791 | 1.174 | 5.201 | ENSMUSG00000039943 | PLCB4 | phospholipas | Cytoplasm | enzyme |
| 0.0121 | -1.289 | -0.253 | ENSMUSG00000086233 | Gm11816 |  | Other | other |
| 0.0141 | -0.416 | 5.796 | ENSMUSG00000025369 | SMARCC2 | SWI/SNF rela | Nucleus | transcription regulator |
| 0.0141 | -0.817 | 1.858 | ENSMUSG00000041961 | ZNRF3 | zinc and ring | Plasma Memk | enzyme |
| 0.0259 | 0.744 | 3.852 | ENSMUSG00000027692 | TNIK | TRAF2 and N | Plasma Memk | kinase |
| 0.0259 | 0.788 | 2.593 | ENSMUSG00000024563 | SMAD2 | SMAD family | Nucleus | transcription regulator |
| 0.0266 | -2.594 | 1.017 | ENSMUSG00000098022 | ZFP82 | ZFP82 zinc fi | Nucleus | transcription regulator |
| 0.0274 | -1.197 | 7.892 | ENSMUSG00000025809 | ITGB1 | integrin subu | Plasma Memk | transmembrane receptor |
| 0.0303 | -2.182 | 2.224 | ENSMUSG00000029863 | CASP2 | caspase 2 | Cytoplasm | peptidase |
| 0.0303 | -0.743 | 1.824 | ENSMUSG00000040747 | CD53 | CD53 molecu | Plasma Memk | other |
| 0.0303 | 1.474 | 3.733 | ENSMUSG00000030532 | HDDC3 | HD domain co | Other | other |
| 0.0303 | 1.285 | 4.582 | ENSMUSG00000022257 | LAPTM4B | lysosomal pro | Cytoplasm | other |
| 0.0303 | -1.197 | 5.872 | ENSMUSG00000035545 | LENG8 | leukocyte rec | Other | other |
| 0.0303 | 1.123 | 4.33 | ENSMUSG00000021518 | PTDSS1 | phosphatidyls | Cytoplasm | enzyme |
| 0.0303 | 1.512 | 6.343 | ENSMUSG00000039844 | RAPGEF1 | Rap guanine | Cytoplasm | other |
| 0.0303 | -0.866 | 1.482 | ENSMUSG00000103651 | Gm37206 |  | Other | other |
| 0.0303 | -0.914 | 4.396 | ENSMUSG00000053931 | CNN3 | calponin 3 | Cytoplasm | other |
| 0.0303 | -0.602 | 2.74 | ENSMUSG00000031239 | ITM2A | integral memk | Plasma Memk | other |
| 0.0303 | 0.464 | 4.206 | ENSMUSG00000026047 | POGLUT2 | protein O-gluc | Cytoplasm | enzyme |
| 0.0303 | 1.616 | 5.365 | ENSMUSG00000030861 | ACADSB | acyl-CoA deh | Cytoplasm | enzyme |
| 0.031 | -1.155 | 4.395 | ENSMUSG00000032724 | ABTB2 | ankyrin repea | Nucleus | other |
| 0.031 | -1.41 | 3.626 | ENSMUSG00000059439 | BCAS3 | BCAS3 micro | Nucleus | other |
| 0.031 | -1.329 | 2.405 | ENSMUSG00000008090 | FGFRL1 | fibroblast gro | Plasma Memk | transmembrane receptor |
| 0.031 | 1.648 | 3.078 | ENSMUSG00000037692 | AHDC1 | AT-hook DNA | Nucleus | transcription regulator |
| 0.031 | 0.843 | 4.987 | ENSMUSG00000029647 | PAN3 | poly(A) specif | Cytoplasm | enzyme |
| 0.0362 | -0.417 | 3.396 | ENSMUSG00000081272 | Ap2m1-ps | adaptor-relate | Other | other |
| 0.0391 | -1.994 | 1.382 | ENSMUSG00000106296 | 4632404M16 | RIKEN cDNA | Other | other |
| 0.0391 | -0.944 | 5.655 | ENSMUSG00000053907 | MAT2A | methionine ac | Cytoplasm | enzyme |

## Notes

### Competing Interest Statement

The authors have declared no competing interest.

## REFERENCES

1. Desai N, Olewinska E, Famulska A, Remuzat C, Francois C, Folkerts K. Heart failure with mildly reduced and preserved ejection fraction: A review of disease burden and remaining unmet medical needs within a new treatment landscape. Heart Fail Rev. 2024;29:631–662.

2. Tsao CW, Lyass A, Enserro D, Larson MG, Ho JE, Kizer JR, Gottdiener JS, Psaty BM, Vasan RS. Temporal trends in the incidence of and mortality associated with heart failure with preserved and reduced ejection fraction. JACC Heart Fail. 2018;6:678–685.

3. Dunlay SM, Roger VL, Redfield MM. Epidemiology of heart failure with preserved ejection fraction. Nat Rev Cardiol. 2017;14:591–602.

4. Omote K, Verbrugge FH, Borlaug BA. Heart failure with preserved ejection fraction: mechanisms and treatment strategies. Annu Rev Med. 2022;73:321– 337.

5. Mishra S, Kass DA. Cellular and molecular pathobiology of heart failure with preserved ejection fraction. Nat Rev Cardiol. 2021;18:400–423.

6. McCarroll CS, He W, Foote K, Bradley A, McGlynn K, Vidler F, Nixon C, Nather K, Fattah C, Riddell A, Bowman P, Elliott EB, Bell M, Hawksby C, MacKenzie SM, Morrison LJ, Terry A, Blyth K, Smith GL, McBride MW, Kubin T, Braun T, Nicklin SA, Cameron ER, Loughrey CM. Runx1 deficiency protects against adverse cardiac remodeling after myocardial infarction. Circulation. 2018;137:57–70.

7. Riddell A, McBride M, Braun T, Nicklin SA, Cameron E, Loughrey CM, Martin TP. RUNX1: An emerging therapeutic target for cardiovascular disease. Cardiovasc Res. 2020;116:1410–1423.

8. Li X, Zhang S, Wa M, Liu Z, Hu S. MicroRNA-101 Protects Against Cardiac Remodeling Following Myocardial Infarction via Downregulation of Runt-Related Transcription Factor 1. J Am Heart Assoc. 2019;8:1–16.

9. Zhang D, Liang C, Li P, Yang L, Hao Z, Kong L, Tian X, Guo C, Dong J, Zhang Y, Du B. Runt-related transcription factor 1 (Runx1) aggravates pathological cardiac hypertrophy by promoting p53 expression. J Cell Mol Med. 2021;25:7867–7877.

10. Li P, Jia XY. MicroRNA-18-5p inhibits the oxidative stress and apoptosis of myocardium induced by hypoxia by targeting RUNX1. Eur Rev Med Pharmacol Sci. 2022;26:432–439.

11. Tzimas C, Rau CD, Buergisser PE, Jean-Louis G, Lee K, Chukwuneke J, Dun W, Wang Y, Tsai EJ. WIPI1 is a conserved mediator of right ventricular failure. JCI Insight. 2019;5.

12. Yamaguchi T, Sumida TS, Nomura S, Satoh M, Higo T, Ito M, Ko T, Fujita K, Sweet ME, Sanbe A, Yoshimi K, Manabe I, Sasaoka T, Taylor MRG, Toko H, Takimoto E, Naito AT, Komuro I. Cardiac dopamine D1 receptor triggers ventricular arrhythmia in chronic heart failure. Nat Commun. 2020;11:4364.

13. Yang G, Chen S, Ma A, Lu J, Wang T. Identification of the difference in the pathogenesis in heart failure arising from different etiologies using a microarray dataset. Clinics (Sao Paulo). 2017;72:600–608.

14. Tan WLW, Anene-Nzelu CG, Wong E, Lee CJM, Tan HS, Tang SJ, Perrin A, Wu KX, Zheng W, Ashburn RJ, Pan B, Lee MY, Autio MI, Morley MP, Tam WL, Cheung C, Margulies KB, Chen L, Cappola TP, Loh M, Chambers J, Prabhakar S, Foo RSY, CHARGE-Heart Failure Working Group C-EC. Epigenomes of Human Hearts Reveal New Genetic Variants Relevant for Cardiac Disease and Phenotype. Circ Res. 2020;127:761–777.

15. Sun N, Yazawa M, Liu J, Han L, Sanchez-Freire V, Abilez OJ, Navarrete EG, Hu S, Wang L, Lee A, Pavlovic A, Lin S, Chen R, Hajjar RJ, Snyder MP, Dolmetsch RE, Butte MJ, Ashley EA, Longaker MT, Robbins RC, Wu JC. Patient-specific induced pluripotent stem cells as a model for familial dilated cardiomyopathy. Sci Transl Med. 2012;4:130ra47.

16. Perestrelo AR, Silva AC, Oliver-De La Cruz J, Martino F, Horváth V, Caluori G, Polanský O, Vinarský V, Azzato G, De Marco G, Žampachová V, Skládal P, Pagliari S, Rainer A, Pinto-Do-Ó P, Caravella A, Koci K, Nascimento DS, Forte G. Multiscale Analysis of Extracellular Matrix Remodeling in the Failing Heart. Circ Res. 2021;128:24–38.

17. Edgar R, Domrachev M, Lash AE. Gene Expression Omnibus: NCBI gene expression and hybridization array data repository. Nucleic Acids Res. 2002;30:207–10.

18. Zile MR, Baicu CF, Ikonomidis JS, Stroud RE, Nietert PJ, Bradshaw AD, Slater R, Palmer BM, Van Buren P, Meyer M, Redfield MM, Bull DA, Granzier HL, LeWinter MM. Myocardial stiffness in patients with heart failure and a preserved ejection fraction: contributions of collagen and titin. Circulation. 2015;131:1247–59.

19. Schiattarella GG, Altamirano F, Tong D, French KM, Villalobos E, Kim SY, Luo X, Jiang N, May HI, Wang Z V., Hill TM, Mammen PPA, Huang J, Lee DI, Hahn VS, Sharma K, Kass DA, Lavandero S, Gillette TG, Hill JA. Nitrosative stress drives heart failure with preserved ejection fraction. Nature. 2019;568:351–356.

20. Martin TP, MacDonald EA, Bradley A, Watson H, Saxena P, Rog-Zielinska EA, Raheem A, Fisher S, Elbassioni AAM, Almuzaini O, Booth C, Campbell M, Riddell A, Herzyk P, Blyth K, Nixon C, Zentilin L, Berry C, Braun T, Giacca M, McBride MW, Nicklin SA, Cameron ER, Loughrey CM. Ribonucleicacid interference or small molecule inhibition of Runx 1 in the border zone prevents cardiac contractile dysfunction following myocardial infarction. Cardiovasc Res [Internet]. 2023;1–9. Available from: https://academic.oup.com/cardiovascres/advance-article/doi/10.1093/cvr/cvad107/7222852

21. Tong D, Schiattarella GG, Jiang N, May HI, Lavandero S, Gillette TG, Hill JA. Female Sex Is Protective in a Preclinical Model of Heart Failure with Preserved Ejection Fraction. Circulation. 2019;140:1769–1771.

22. Matsiukevich D, Kovacs A, Li T, Kokkonen-Simon K, Matkovich SJ, Oladipupo SS, Ornitz DM. Characterization of a robust mouse model of heart failure with preserved ejection fraction. Am J Physiol Heart Circ Physiol [Internet]. 2023;325:H203–H231. Available from: http://www.ncbi.nlm.nih.gov/pubmed/37204871

23. Liu X, Yin K, Chen L, Chen W, Li W, Zhang T, Sun Y, Yuan M, Wang H, Song Y, Wang S, Hu S, Zhou Z. Lineage-specific regulatory changes in hypertrophic cardiomyopathy unraveled by single-nucleus RNA-seq and spatial transcriptomics. Cell Discov [Internet]. 2023;9:6. Available from: http://www.ncbi.nlm.nih.gov/pubmed/36646705

24. Ni T, Huang X, Pan S, Lu Z. Dihydrolycorine Attenuates Cardiac Fibrosis and Dysfunction by Downregulating Runx1 following Myocardial Infarction. Oxid Med Cell Longev. 2021;2021.

